# *In vivo* risk assessment of yellow fever virus transmission through breastfeeding, and mechanistic insights

**DOI:** 10.1101/2025.07.23.666308

**Authors:** Jeanne Pascard, Sophie Desgraupes, Aurelie Chiche, Patricia Jeannin, Rebecca Kanaan, Antoine Gessain, Han Li, Pierre-Emmanuel Ceccaldi, Aurore Vidy

## Abstract

Yellow fever virus (YFV), a mosquito-borne flavivirus, remains a significant public health threat, especially in areas with low vaccine coverage. Since 2010, yellowfevervaccination is not recommended for breastfeeding women due to reported cases of vaccine strain transmission through breast milk, leading to neonatal meningoencephalitis. However, the efficiency of YFV vaccine strain transmission via breastfeeding remains unknown, and wild-type strains transmission may be suspected based on viral RNA detection in breast milk. Direct evidence of breastfeeding-related transmission in humans is challenging to obtain given the confounding presence of vector-borne transmission, making animal models crucial for evaluating this risk. In this study, the A129 mouse model was used to investigate YFV transmission via breastfeeding for both wild-type and vaccine strains. Results show that both strains can spread to mammary glands, leading to viral detection in breast milk as free viral particles and cell-associated virus, with similar viral loads for all strains. Mammary stromal and immune cells are primary targets of YFV *in vivo,* while mammary epithelial cells also support infection, suggesting two possible mechanisms of mammary epithelial crossing. Neonates are found to be susceptible to oral infection, with higher infection rates for the wild-type strain but evidence of neuroinvasion for both strains. Both strains can infect and cross an *in vitro* human intestinal barrier model, indicating this epithelium as a potential viral entry site for neonates. Finally, this study confirms the existence of YFV transmission through breastfeeding in an animal model, highlighting the need to consider it among transmission risks.

## INTRODUCTION

Yellow fever virus (YFV) is a mosquito-borne orthoflavivirus that causes a potentially fatal haemorrhagic disease, with ∼200,000 cases and 30,000 deaths annually worldwide^1^. Clinical manifestations range from mild symptoms to severe jaundice and haemorrhagic fever, with fatality rates up to 60% in severe cases^2^• Despite the availability of a safe and effective live-attenuated vaccine (17D), YFV remains endemic in 44 countries, and outbreaks continue due to insufficient immunization coverage^3^• Two major vaccine substrains are in global use: 17D-204 and 17DD^4^•

In addition to mosquito-borne transmission, YFV has been implicated in non-vectorial routes such as blood transfusion^5,6^ organ transplantation^6^, and vertical transmission^7–12^. Notably, several cases of breastfeeding transmission involving YFV vaccine strains have been reported. In each instance, exclusively breastfed infants developed meningoencephalitis shortly after maternal vaccination^9–12^. Detection of vaccine RNA or YFV-specific lgM in the infants’ serum and/or cerebrospinal fluid provided compelling evidence of transmission via breastfeeding^9 11^ As a result, since 2010, the U.S. Centers for Disease Control and Prevention (CDC) has recommended against vaccinating breastfeeding women, except for epidemics or unavoidable travel to endemic areas ^3^

Although less documented, wild-type YFV transmission via breastfeeding remains biologically plausible. During Brazil’s 2016-2018 outbreak, wild-type YFV RNA was detected in breast milk of an infected mother whose infant also developed symptoms^14^- raising concern about this potential, but often underestimated, route of transmission. However, establishing breastfeeding transmission of mosquito-borne viruses in humans is particularly difficult in endemic areas, where the omnipresence of vectors makes it nearly impossible to exclude mosquito-borne transmission.

To address this challenge, criteria adapted from Koch’s postulates have been proposed^15^. These include epidemiological evidence of breastfed infant infection, detection of infectious virus in breast milk, exclusion of alternative transmission routes, and reproduction of transmission via oral or breastfeeding routes in animal models. While epidemiological data are informative, animal models are essential to confirm causality in a controlled environment. Importantly, transmission of the YFV 17D vaccine strain through breastfeeding is considered proven, as this attenuated strain cannot be transmitted by mosquitoes^16^-thereby ruling out vectorial transmission and strengthening the causal link.

Using these criteria, breastfeeding transmission has been confirmed for three human viruses: human T-cell lymphotropic virus type 1 (HTLV-1), human immunodeficiency virus (HIV), and human cytomegalovirus (CMV)^15^. It is also suspected for several arboviruses. Infant infections have been reported following maternal infection with chikungunya virus (CHIKV), dengue virus (DENV), West Nile virus (WNV), and Zika virus (ZIKV), with viral genomes from all four-and infectious DENV and ZIKV particles-detected in breast milk^17^•

In this study, we used *in vivo* (mouse) and *in vitro* (human cells) models to investigate YFV transmission to neonates via oral exposure and breastfeeding. We demonstrate that both YFV vaccine and wild-type strains can infect the mammary gland, cross the mammary epithelium, and are excreted in murine milk as infectious free particles and cell­ associated virus. *In vivo,* YFV primarily targets mammary stromal and immune cells, however human mammary epithelial cells are also permissive to infection, suggesting two mechanisms for epithelial crossing in mammary glands: infected immune cell transmigration, so called as "Trojan horse" strategy, and epithelial infection with viral release. We also show that neonatal mice are susceptible to YFV via the oral route, with neuroinvasion following intragastric exposure. Using a human intestinal epithelium model, we demonstrate that both strains can infect and cross this epithelial barrier without disrupting its integrity. Finally, we confirm that YFV can be transmitted to suckling pups via breastfeeding-albeit at a low frequency-, validating this route of transmission for YFV.

## MATERIALS AND METHODS

### Animal model

Experiments used A 129 mouse model, deficient interferon-a/13 receptor (IFNAR-J-). Mice were housed and bred in the lnstitut Pasteur (Paris, France), in animal facilities accredited by the French Ministry of Agriculture for breeding and performing experiments on live rodents (authorization no. A75150101).

### Virus strains

Three YFV strains were used: wild-type strains Asibi (GenBank: AY640589.1) and Dakar/HD1279 (GenBank: MN106242.1), and vaccine strain 170-204 (GenBank: MN708488.1).

### Mouse infection

Non-lactating females were subcutaneously inoculated with 1-3 x 10^5^ plaque-forming units (PFU) of YFV (Asibi, Dakar/HD1279 or 170-204 strains). Blood samples were collected at various days post-inoculation (dpi), and mammary glands were harvested at selected dpi. Lactating females received subcutaneously 1-3 x 10^5^ of YFV (Asibi or 170- 204 strains). Blood samples were collected between 4 and 6 dpi. At 5 or 6 dpi, breast milk and mammary glands were collected^18^. For experiments of breastfeeding-mediated transmission, blood and organ samples were collected from suckling pups between 7 and 9 dpi. For oral infection of neonatal mice, A129 pups were inoculated with YFV (2 x 10^4^ to 1 x 10^5^ PFU, Asibi or 170-204) by intragastric route. Blood samples and brains were collected at 6 dpi. Protocols and animal monitoring are detailed in supplementary materials.

### Ethics statement

Animal experiments complied with French and European regulations on care and protection of laboratory animals (EC Directive 2010/63, French Law 2013-118, 6 February 2013). All experiments were approved by the Ethics Committee n°89 and registered by the French "Ministere de l’Enseignement Superieur, de la Recherche et de l’lnnovation" (MESRI) under the reference "APAFIS#31829-2021052812549296" (date of approval: 31 May 2021). Use of genetically modified mice (A129) was approved by the institutional instances and the MESRI under the reference n°9319 (date of approval: 6 December 2021).

### Statistical ahnalysis

Statistical analyses were performed with GraphPad Prism; statistical tests and replicate numbers are detailed in figure legends. Analyses were supported by the lnstitut Pasteur Bioinformatics and Biostatistics Hub.

***Full protocols and reagent information are available in the supplementary materials*.**

## RESULTS

### YFV disseminates to the mammary glands of non-lactating and lactating A129 mice

To investigate the early steps of YFV transmission via breastfeeding, A129 mice-a well­ established model of YFV infection^19^- were used to assess viral dissemination to mammary glands and subsequent excretion into breast milk. Infection kinetics were first evaluated in non-lactating females for ethical and practical reasons, with key findings later validated in lactating mice.

Mice were subcutaneously inoculated with wild-type Asibi or Dakar/HD1279 strains, or vaccine strain 17D-204 (1-3x 10^5^ PFU), modelling a local infection as occurs following mosquito bite. Clinical signs, weight, and viremia were monitored, and mammary glands collected at various days post-infection (dpi) **(Figure 1A).** No mortality was observed. Only wild-type strains induced mild symptoms and limited weight loss (∼5% for Dakar), while infection with 17D-204 strain remained asymptomatic **(Supplementary Figure S1A-B).** Viremia was confirmed in all groups by the presence of viral RNA (vRNA) in the plasma, except in three 17D-204 cases, where infection was confirmed in the spleen **(Figure S1C).**

**Figure 1:**
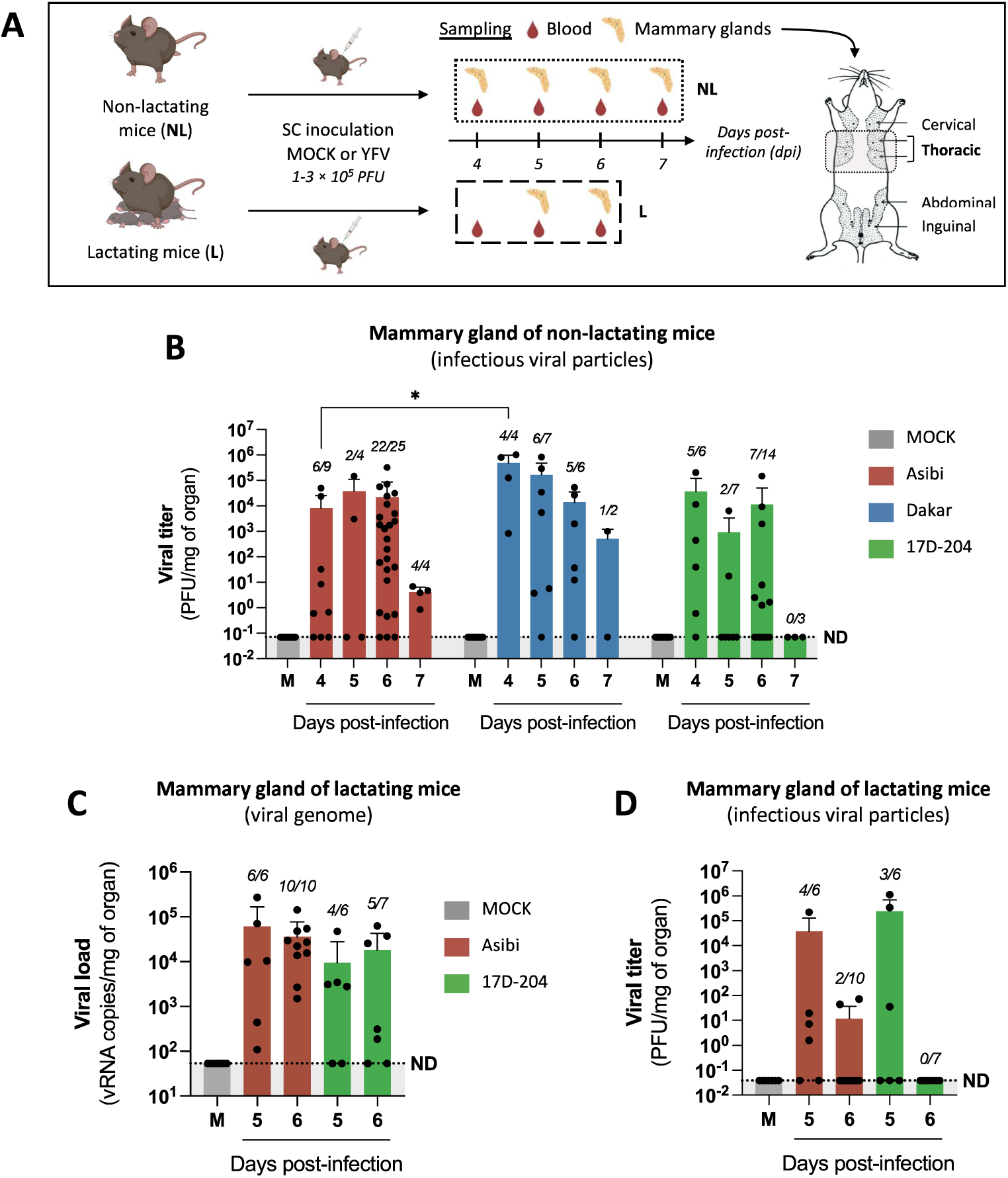
YFV disseminates to the mammary glands of non-lactating and lactating A129 mice. **(A)** Non-lactating (6- to 23-week-old) and lactating (10- to 22-week-old) A129 mice were either MOCK-inoculated or inoculated with YFV via subcutaneous (SC) injection. Non­ lactating females were inoculated with PBS (MOCK, n=16) or with 1 to 3 x 1as PFU of the Asibi (n=42), Dakar/HD1279 (n=19) or 17D-204 strains (n=30) of YFV (7 independent experiments). Lactating females were inoculated 1 to 13 days post-partum with PBS (MOCK, n=7) or with 1 to 3 x 1as PFU of the Asibi (n=16) or 17D-204 strains (n=13) (5 independent experiments). At various days post-infection (dpi), blood samples and thoracic mammary glands were collected to assess mice infection and mammary gland dissemination, respectively (Illustration created with Biorender). **(B)** YFV dissemination to thoracic mammary glands of non-lactating mice was assessed by plaque assay on Vero cells to detect infectious viral particles. **(C-D)** Viral dissemination to thoracic mammary glands of lactating mice was assessed by RT-qPCR using NS3-specific primers to determine the presence of viral RNA genome (C) and by plaque assay on Vero cells (D). *The number of infected mammary glands relative to the total number of tested mammary glands is shown in italics above each time point.* Results are expressed as the mean ± standard deviation. The dashed lines indicate the specificity limit, which represents the threshold under which values were considered as "not detected" (ND). Statistical tests: Kruskal-Wallis test with Dunn’s multiple comparisons was used all in panels, comparing means values between all strains at all time-points. * = p < 0.05. Only statistically significant differences (p < 0.05) are indicated.

In non-lactating mice, infectious virus was detected by plaque assay in mammary glands for all strains, peaking between 4-6 dpi with notable inter-individual variation **(Figure 1B).** Titers declined by 7 dpi, suggesting limited persistence in this tissue. When pooling all time points, mammary infection was significantly more frequent with wild-type Asibi (73.8%) and Dakar (84.2%) than with 17D-204 (46.7%) **(Figure S1D, a),** suggesting limited dissemination of the vaccine strain.

Using the identified peak infection times (5-6 dpi), presence of vRNA and infectious particles in mammary glands of lactating females was confirmed **(Figure 1C-D),** with marked inter-individual variability. RT-qPCR detection was significantly more frequent in Asibi-infected mice (100%) than in 17D-204-infected ones (69.2%) **(Figure S1D, b),** further supporting reduced dissemination of the vaccine strain. Notably, infectious virus was detected less frequently in lactating than in non-lactating glands, likely due to technical challenges in viral quantification, as milk and tissue remodelling may reduce detection sensitivity.

These results show that both wild-type and vaccine YFV strains can reach the mammary glands in mice under both lactating and non-lactating conditions.

### YFV infectious virus is released into murine breast milk, as free viral particles and cell-associated virus

To investigate viral excretion into milk, lactatingA129 mice were inoculated with YFV Asibi or 17D-204 strains via subcutaneous route **(Figure 2A),** and successful infection was confirmed in plasma or spleen **(Figure S2A).** Viral RNA was detected in breast milk collected at 5-6 dpi for both strains in 80-90% of animals **(Figure S2C, a),** ranging from 2 x 10^3^ to 7.1 x 1as (Asibi) and 2.5 x 10^4^ to 4.7 x 10^6^ vRNA copies/ml (17D-204) **(Figure S2B, a),** reflecting a high rate of viral shedding into milk. Infectious virus was also detected, with higher titers at 5 dpi - generally greater for 17D-204 (up to 1.5 x 1as PFU/ml) than Asibi (up to 3x 10^4^ PFU/ml)-though differences were not statistically significant **(Figure 2B).** Titers declined by 6 dpi, with infectious virus undetectable in the Asibi group and lower detection in 17D-204 samples.

**Figure 2:**
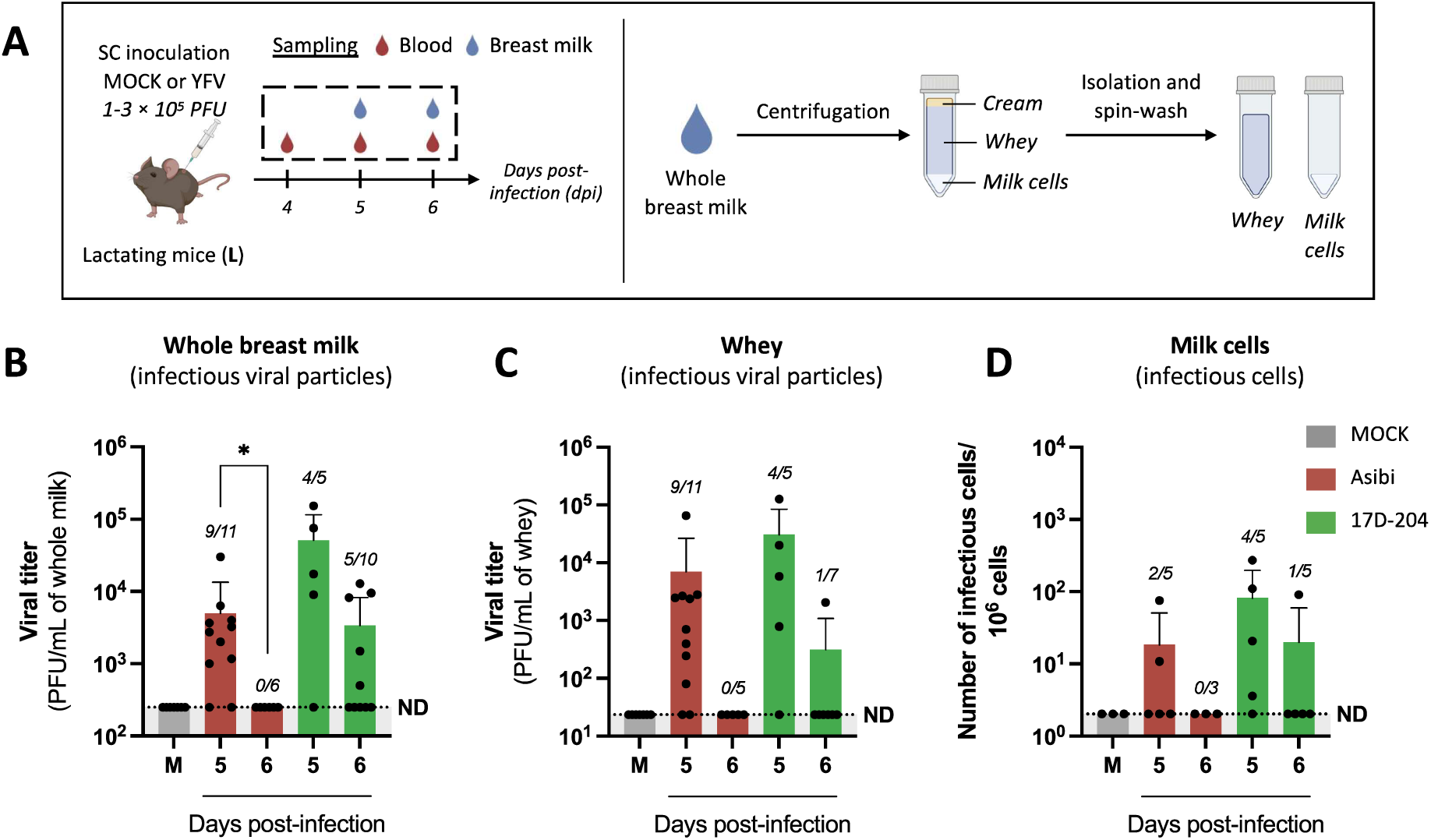
YFV infectious virus is released into murine breast milk, both as free viral particles and cell-associated virus. **(A)** Lactating A129 mice (10-22 weeks old) were either MOCK-inoculated (n=7) or inoculated with YFV at 1 to 12 days post-partum via subcutaneous (SC) injection of 1 to 3 x 1as PFU of the Asibi (n=17) or 17D-204 (n=15) strains of YFV (6 independent experiments). Four to 6 days after inoculation (dpi), blood and breast milk samples were collected. For some milk samples, a portion was centrifuged to separate cream, whey, and cells. Whey was further clarified by three centrifugation to enhance purity, while the cell pellet underwent three wash-spin cycles in PBS to remove free viral particles (Illustration created with Biorender). **(B-C)** Detection of YFV in whole breast milk (B) and whey (C) was assessed by plaque assay on Vero cells to determine the presence of infectious viral particles. **(D)** Detection of infectious cell-associated virus in the milk cell pellet was performed by infectious center assay. *The number of infected samples relative to the total number of tested samples per condition* is *shown in italics above each time point.* Results are expressed as the mean± standard deviation. The dashed lines indicate the specificity limit, which represents the threshold under which values were considered as "not detected" (ND). Statistical tests: Kruskal-Wallis test with Dunn’s multiple comparisons was used in all panels, comparing means values between all strains at all time-points.*= p < 0.05. Only statistically significant differences (p < 0.05) are indicated.

To characterize the form of virus in milk, murine milk samples were fractionated by centrifugation into cream, whey and cellular components **(Figure 2A).** vRNA was consistently detected in whey and purified milk cells **(Figure S2B, a-b),** and infectious virus was found in whey **(Figure 2C)** and in milk cells after co-culture with permissive cells **(Figure 2D),** indicating the presence of both free viral particles and cell-associated infectious virus in breast milk.

These findings demonstrate that both wild-type and vaccine YFV strains are excreted into breast milk as free and cell-associated infectious virus. No strain-specific differences were observed in viral load, infectious titer, or the proportion of positive milk samples **(Figure S2C),** which contrasts with the dissemination patterns seen in mammary tissue. This viral excretion into breast milk requires crossing the mammary epithelium, via mechanisms yet to be elucidated.

### YFV infects mainly immune and stromal cells in the mammary glands of non­ lactating mice *in vivo*

The mammary gland comprises epithelial structures within a fibro-adipose matrix, featuring various cell types. Stromal cells (including adipocytes and fibroblasts) provide structural support, and endothelial and immune cells manage vascularization and immune surveillance^20^• The pseudostratified epithelium includes luminal cells (milk secretion) and contractile myoepithelial cells (milk ejection). YFV presence in breast milk suggests mammary epithelium viral crossing, potentially through infected immune cell transmigration or epithelial cell infection followed by viral release into the milk or shedding of infected cells.

To study YFV tropism, mice were subcutaneously inoculated with YFV (Asibi and Dakar strains). Infection was confirmed in plasma **(Figure S3A),** and mammary glands were analyzed at 6 dpi using enzymatic dissociation and flow cytometry or immunohistochemistry **(Figure 3A).** Mammary cells were categorized into endothelial (CD31+), immune (CD45+), epithelial (CD24+), and stromal cells (CD31-co4s-co24- CD49fl Luminal (CD24highCD49f^10^W) and myoepithelial cells (CD24^10^WCD49fhigh) were further identified within the epithelial subset **(Figure S3B).** Non-lactating mice were selected to obtain the large number of mammary cells necessary for flow cytometry, and to avoid complications from milk presence and gland morphology during immunohistochemistry.

**Figure 3:**
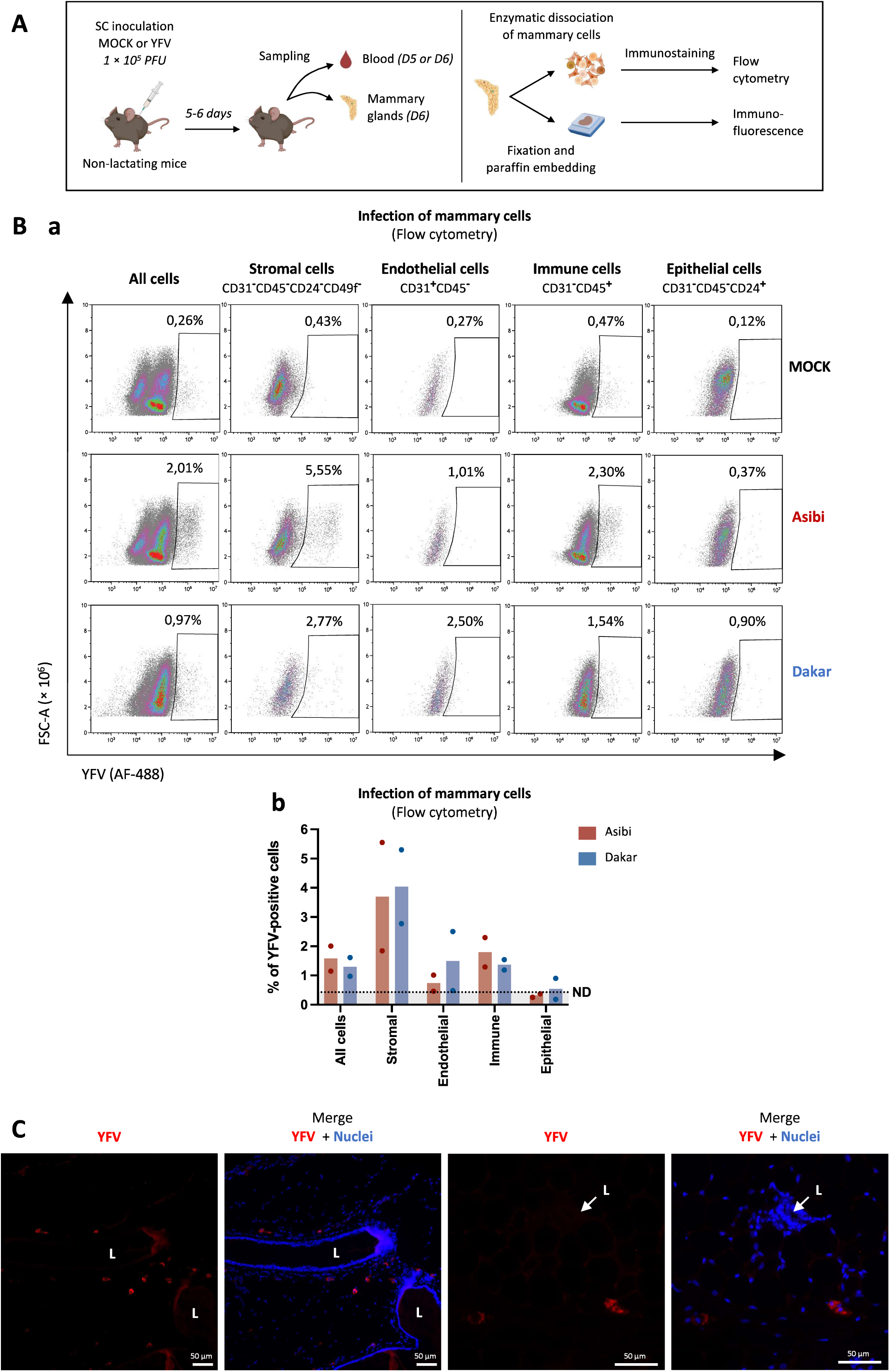
YFV infects mainly immune and stromal cells in the mammary glands of non-lactating mice *in vivo*. **(A)** Non-lactating A129 mice (11-19 weeks old) were either MOCK-inoculated or inoculated with YFV via subcutaneous (SC) injection of 1 x10^5^ PFU of the Asibi or Dakar strains of YFV. At 5-6 days after inoculation (D5-D6), blood and thoracic and/or abdominal mammary samples were collected to assess mice infection and viral tropism in the mammary tissue. Mammary glands were analysed by flow cytometry following enzymatic dissociation and immunostaining, (2 independent experiments; MOCK n=7, Asibi n=7, Dakar n=5) or by immunohistochemistry after fixation, paraffin embedding, sectionning and immunostaining (2 independent experiments; MOCK n=1, Asibi n=4) (Illustration created with Biorender). **(B)** For flow cytometry, thoracic and abdominal mammary glands from 3-4 mice per group were pooled and enzymatically digested. Cell suspensions were stained with fluorochrome-conjugated antibodies against CD31 (PE), CD45 (APC), CD24 (BV421), and CD49f (PECy7), then fixed, permeabilized, and stained for YFV using mouse polyclonal antibodies (ascitic fluid) and an AF488-conjugated secondary antibody. Samples were analyzed by flow cytometry. Panel (a) shows representative plots; panel (b) quantifies YFV+ cells (2 independent experiments, one dot = one experiment). Dashed lines indicate the specificity threshold (ND: not detected). **(C)** For immunohistochemistry, mammary glands from Asibi-infected mice were fixed in 10% formalin, paraffin­ embedded, sectioned (5 µm), and stained for YFV using the same primary antibody and an AF546-conjugated secondary. Fluorescence microscopy was used for analysis. Red= YFV; blue= nuclei (DAPI). L: lumen of lactiferous ducts, lined by mammary epithelium and surrounded by stroma. Both images are from the same animal and are representative of four independent animals.

Flow cytometry revealed YFV infection in a small proportion of mammary cells, predominantly in stromal and immune cells (∼4 and 1.5% of infected cells, respectively), with minimal detection in endothelial and epithelial cells **(Figures 3B, S3C).** lmmunohistochemistry of mammary glands confirmed presence of YFV-positive cells mainly in the stromal compartment, likely infiltrating immune cells or resident stromal cells **(Figure 3C).**

These findings identify mammary stromal and immune cells as possible primary YFV targets *in vivo,* suggesting that infected immune cell transmigration may facilitate viral crossing of the mammary epithelium. Nevertheless, direct epithelial cell infection cannot be excluded, as it might occur below detection thresholds or be influenced by technical limitations, as being among the most difficult cells to dissociate. Given that epithelial cells constitute the mammary barrier and are predominant in human and murine milk (up to 98 and 88% of milk cells, respectively^18•21^), their potential infection, even if rare, warrants further *in vitro* investigation as an alternative mechanism for viral crossing through the mammary epithelium.

### Human primary mammary epithelial cells, as well as luminal and myoepithelial cell lines, are permissive to YFV infection

To evaluate whether productive infection of mammary epithelial cells could facilitate YFV crossing of the mammary barrier, and whether susceptibility varies among epithelial subtypes, human primary mammary epithelial cells (HMEpiC) and two human mammary epithelial cell lines-luminal MCF7 and myoepithelial MDA-MB-231-were exposed to three YFV strains (Asibi, Dakar, and 17D-204) **(Figure 4A).** Viral replication and release were assessed by plaque assay over time.

**Figure 4:**
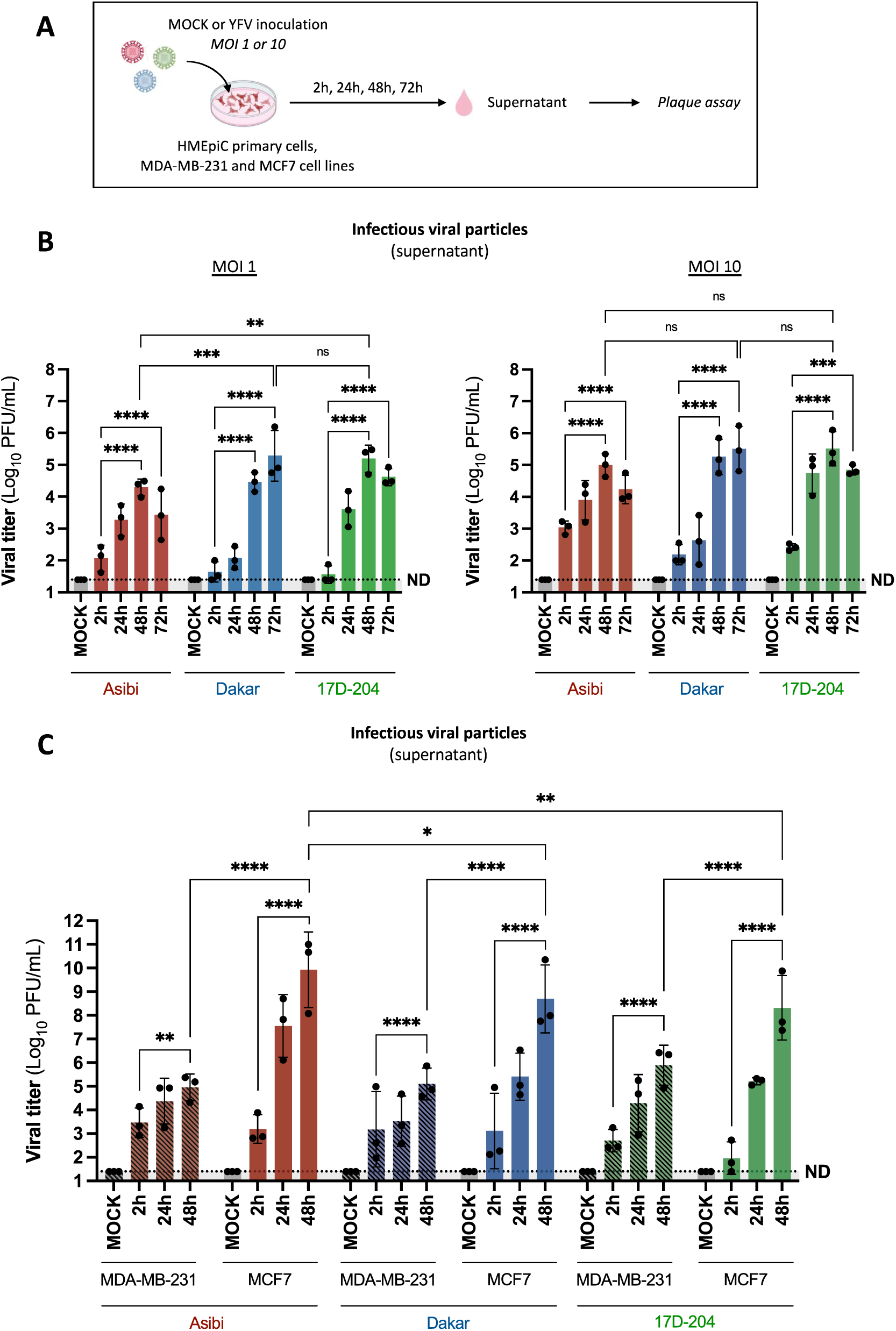
Human primary mammary epithelial cells, as well as luminal and myoepithelial cell lines, are permissive to YFV infection. **(A)** Human primary mammary epithelial cells (HMEpiC), as well as myoepithelial (MDA­ MB-231) and luminal (MCF7) human mammary epithelial cell lines, were exposed to three YFV strains (Asibi, Dakar/HD1279 or 17D-204)- at multiplicities of infection (MOI) of 1 or 10 for HMEpiC, and MOl 1 for MDA-MB-231 and MCF7 cell lines. Control cells were MOCK­ treated. At different times post-inoculation, viral production in supernatants was measured by plaque assay (Illustration created with Biorender). **(B-C)** Culture supernatants were collected, and infectious viral particles quantification was assessed by plaque assay on Vero cells for HMEpiC cells (B; MOI 1 and 10) and for MDA-MB-231 and MCF7 cell lines (C; MOl 1). Results are expressed as the mean± standard deviation. The dashed lines indicate the specificity limit, which represents the threshold under which values were considered as "not detected" (ND). The results in (B) and (C) are representative of three independent experiments; in (B), data correspond to two different cell batches. Dots represent the mean value of each experiment. Statistical test: Ordinary two-way ANOVA followed by Tukey’s multiple comparisons test, performed on log10- transformed data (using all individual data points, not only experiment means), was used in panels (B-C). For statistical analyses: * = p < 0.05; ** = p < 0.005; *** = p < 0.0005; **** = p < 0.0001.

**Figure 5:**
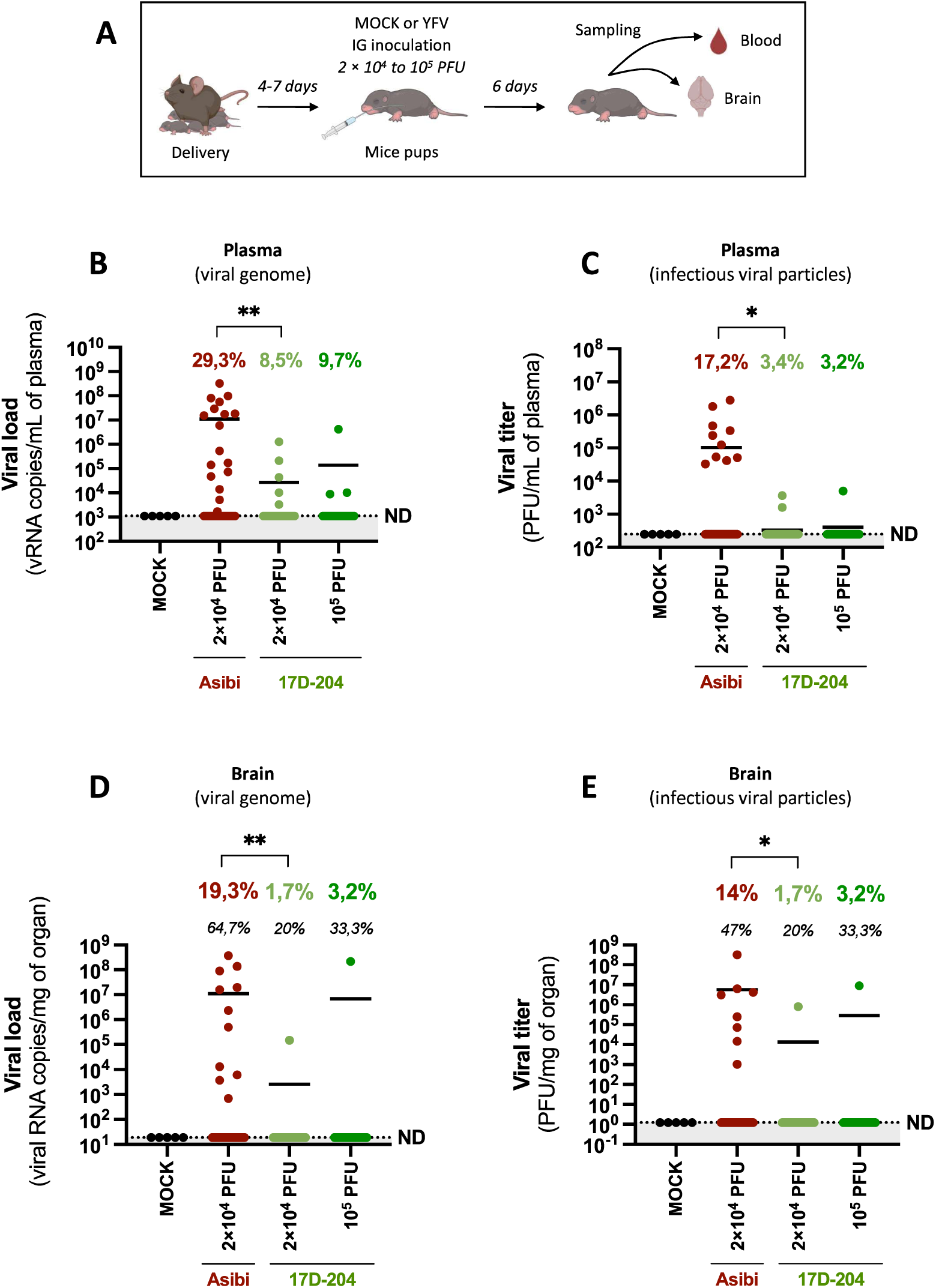
Oral inoculation of A129 mouse pups with YFV results in systemic infection and can lead to neuroinvasion. **(A)** A129 mouse pups (4- to 7-days-old) were inoculated via the intragastric (IG) route with yellow fever virus (YFV) or with PBS as a negative control (MOCK). Pups received either PBS (MOCK, 2 litters, n = 5), 2 x10^4^ PFU of the Asibi strain (10 litters, n = 58), or 2 x10^4^ PFU (11 litters, n = 59) or 1 x10^5^ PFU (4 litters, n = 31) of the 17D-204 strain of YFV (6 independent experiments). Blood and brain samples were collected at 6 days post­ infection to assess viremia (B-C) and brain infection (D-E) (Illustration created with Biorender). **(B-C)** Viral infection in mouse pups was assessed in plasma (obtained by blood centrifugation) using NS3-specific RT-qPCR to quantify viral RNA load (B), and by plaque assay on Vero cells to determine infectious viral titers (C). **(D-E)** YFV presence in brain tissue was evaluated post-homogenization by RT-qPCR for viral RNA detection (D), and by plaque assay on Vero cells to quantify infectious virus (E). Results are presented as mean values; the dashed lines indicate the specificity limit, which represents the threshold under which values were considered as "not detected" (ND). Percentages in bold (B-E) represent the proportion of positive samples out of the total; percentages in italic (D-E) represent the proportion of brain-positive samples among those positive in plasma by RT-qPCR. Statistical test: Fisher’s exact test was used in panels (B-E). For statistical analyses: * = p < 0.05; ** = p < 0.005. Only statistically significant differences (p < 0.05) are indicated.

All cells supported productive infection, with significant time-dependent increases in viral titers for all strains **(Figure 4B-C).** In HMEpiC cells, replication level was dose­ dependent, with higher titers at MOl 1O compared to MOI 1. Peak titers for Asibi and 17D- 204 occurred at 48 hours post-infection (hpi), with 17D-204 showing higher values at MOI 1 (5.19 vs. 3.63 log,_0_ PFU/ml). Dakar titers peaked at 72 hpi (5.29 log_10_ PFU/ml at MOl 1), exceedingAsibi but not significantly different from 17D-204. At MOl 10, peak titers did not differ significantly between strains **(Figure 4B).**

In cell lines, luminal MCF7 cells produced markedly higher viral titers than MDA-MB-231 cells across all strains (e.g., Asibi: 9.92 vs. 4.96-5.89 log,_0_ PFU/ml at 48 hpi). Among the strains, no significant differences were observed in MDA-MB-231 cells, whereas in MCF7 cells, Asibi titers were significantly higher than those of Dakar and 17D-204 **(Figure 4C).** These results indicate that human mammary epithelial cells, both luminal and myoepithelial phenotypes, can support YFV replication, suggesting that the mammary epithelium may contribute to viral excretion into breast milk, either through free virus or infected cell shedding.

### Oral inoculation of A129 mouse pups with YFV results in systemic infection and can lead to neuroinvasion

After excretion in milk, YFV must cross the infant’s digestive barrier (tonsils, intestinal epithelium…) to allow transmission via breastfeeding. To assess potential oral transmission of YFV, susceptibility of mouse pups after intragastric inoculation with wild­ type (Asibi) or vaccine (17D-204) strains (2 x 10^4^ to 1 x 10^5^ PFU) was evaluated. At 6 dpi, blood and brain samples were collected to evaluate systemic infection and neuroinvasion **(Figure SA).**

Plasma vRNA was detected for both strains, with significantly higher infection rates for Asibi (29.3%) than for 17D-204 at either viral dose (8.5% and 9.7%, respectively) **(Figure SB).** Infectious virus was found in pups with high plasma viremia (>5 x 10^5^ vRNA copies/ml), titers being higher for Asibi (up to 2.8 x 10^6^ PFU/ml) than for 17D-204 (up to 3.7 x 10^3^ PFU/ml) **(Figure SC).**

Neuroinvasion occurred for both strains. In Asibi-infected pups, vRNA and infectious virus were detected in the brain of 64.7% and 47% of plasma-positive animals, respectively **(Figure SD-E).** For 17D-204, brain infection was observed in 1/5 and 1/3 plasma-positive pups at 2 x 10^4^ and 1 x 10^5^ PFU, respectively, though the limited sample size precludes conclusions on neuroinvasion efficiency.

These findings demonstrate that oral YFV inoculation can lead to systemic infection and neuroinvasion in some pups, supporting the plausibility of breastfeeding-associated transmission and neurological complications observed in human cases involving YFV vaccine strains^9–12•^

### YFV can infect and cross an *in vitro* model of human intestinal epithelium without disrupting its integrity

Oral infection suggests that YFV can cross a digestive barrier, such as the tonsillar or intestinal mucosa. To investigate whether the intestinal epithelium could allow YFV entry following oral exposure, a previously established *in vitro* model of human intestinal epithelium based on Caco-2/TC7 cells cultured on Transwell inserts was used. After 14-29 days, tight polarized monolayers (transepithelial electrical resistance (TEER) >250 O.cm^2^) were apically inoculated at MOl 1 with YFV (Asibi, Dakar, 17D-204 strains). Barrier integrity was monitored by TEER measurement and tight junction protein ZO-1 immunostaining. Intracellular RT-qPCR and plaque assays of apical and basolateral media were used to assess viral replication, production, and epithelial crossing **(Figure 6A).**

**Figure 6:**
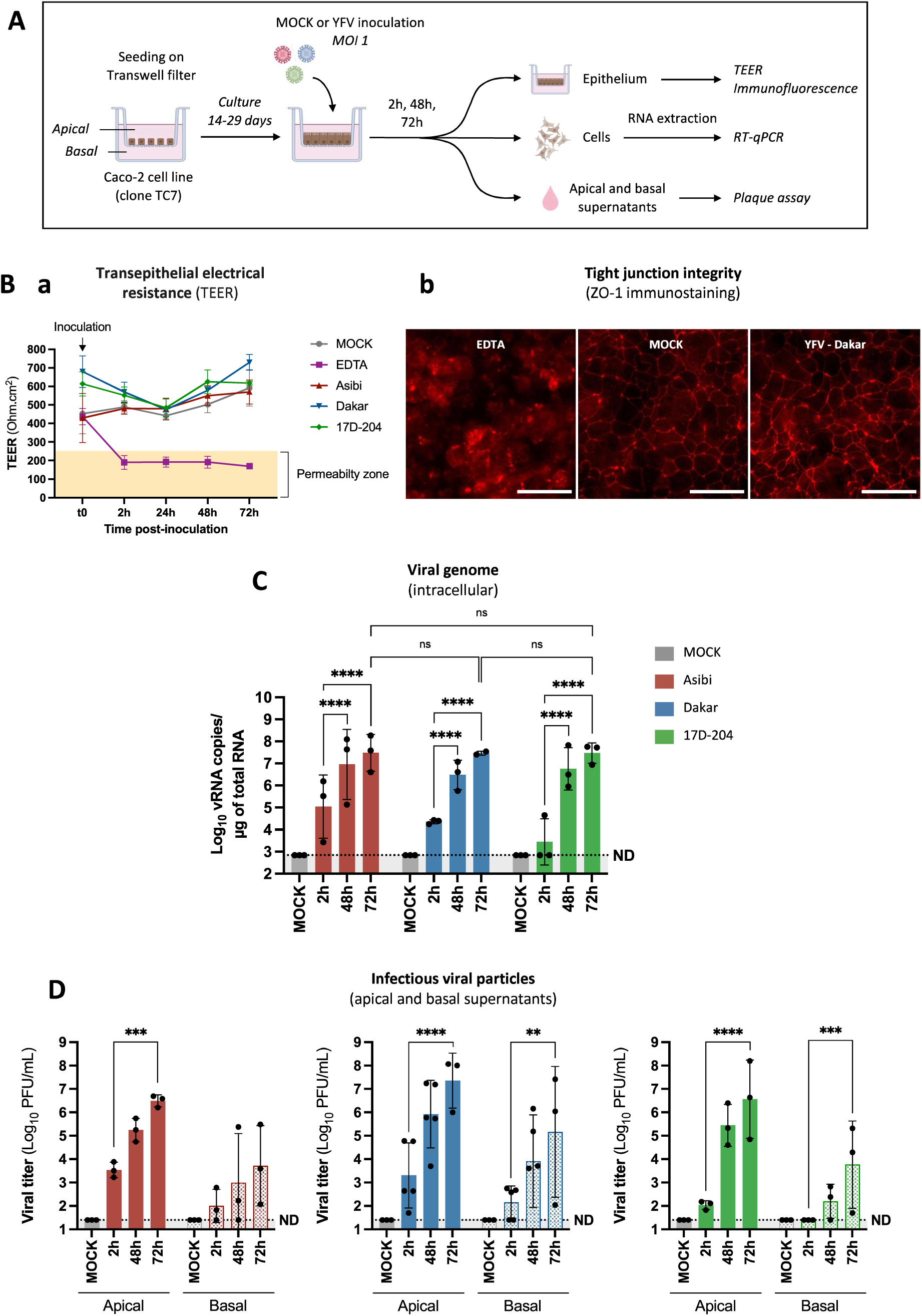
YFV can infect and cross an *in vitro* model of human intestinal epithelium without disrupting its integrity. **(A)** Caco-2/TC7 cells (3 x 10^4^) were seeded into the upper chamber of Transwell inserts (12 mm diameter, 3 µm pores) and cultured for 14-29 days to form a tight, polarized intestinal epithelial monolayer separating apical (upper) and basolateral (lower) compartments. Monolayer integrity was assessed by transepithelial electrical resistance (TEER) measurements. Cells were apically infected with YFV strains (Asibi, Dakar/HD1279 or 17D-204) at MOI 1, or left uninfected as controls (MOCK). At various time points post-infection, TEER and ZO-1 immunostaining were used to monitor barrier integrity (B), viral replication was quantified by RT-qPCR on intracellular RNA (C), and viral production and crossing of the epithelium was assessed by plaque assay on apical and basolateral supernatants (D) (Illustration created with Biorender). **(B)(a)** TEER values were monitored over time. EDTA (12.5 mM; purple) was used as a positive control to disrupt barrier function. The yellow zone (0-250 Ohm.cm^2^) indicates the permeability threshold. Data shown are representative of two independent experiments and consistent across all replicates. (b) At 48 h post-infection, monolayer morphology was visualized by immunofluorescence of the tight junction protein ZO-1 (red). Scale bar: 50 µm. **(C)** Viral RNA levels were measured at 2h, 48h, and 72h post-infection. RNA was extracted from Caco-2/TC7 cells, and YFV RNA was quantified using NS3-specific RT-qPCR. Results are expressed as viral RNA copies/µg of total RNA, calculated using a standard curve generated from 10-fold serial dilutions of a quantified YFV NS3 plasmid. **(D)** At 2h, 48h, and 72h post-infection, apical and basolateral supernatants were collected and infectious virus quantified by plaque assay on Vero cells. Results in panels (A), (C), and (D) are expressed as mean ± standard deviation. Dashed lines represent the detection threshold, below which values were considered "not detected" (ND). Data in panels (C) and (D) are representative of minimum three independent experiments, with each dot corresponding to the mean value from one experiment. Statistical test: Ordinary two-way ANOVA followed by Tukey’s multiple comparisons test, performed on logrn-transformed data, was used in panel (C); Ordinary one-way ANOVA followed by Tukey’s multiple comparisons test, performed separately for strain and apical/basal supernatants and on log,a-transformed data (using all individual data points, not only experiment means), was used in panel (D). For statistical analyses: * = p < 0.05; ** = p < 0.005; *** = p < 0.0005; **** = p < 0.0001.

The TEER values remained stable in YFV-infected monolayers, in contrast to EDTA-treated controls, which showed a marked decrease **(Figure 6B, a).** ZO-1 immunoreactivity at 48 hpi showed intacttightjunction distribution **(Figure 6B, b),** confirming a preserved barrier integrity.

Intracellular vRNA increased significantly over time for all strains, peaking at 72 hpi (∼7.47 log_10_ copies/µg) and confirming YFV replication in differentiated enterocytes **(Figure SC).** Infectious titers showed significant increase apically and basolaterally for all strains, though the increase for Asibi on the basolateral side was not statistically significant **(Figure 6D).** Basolateral titers reached high levels by 72 hpi (Asibi: 3.71, Dakar: 5.17, 17D- 204: 3.77 log_10_ PFU/ml), supporting productive infection in the basolateral compartment, though passive transcytosis of viral particles cannot be excluded.

These results supportthe intestinal epithelium as a potential portal of entry for both wild­ type and vaccine YFV strains via productive infection of enterocytes.

### Limited but detectable transmission of YFV to suckling pups through breastfeeding

After validating the sequential steps of milk-borne YFV transmission, breastfeeding transmission experiments were performed in our mouse model to confirm this route of infection.

Lactating A129 mice were inoculated post-partum with YFV (Asibi and 17D-204 strains) at 3 x 10^5^ PFU. At 7 and/or 9 dpi, samples from suckling pups (blood, spleen, liver) were collected to assess infection **(Figure 7A).** Dams’ infection was confirmed by viremia **(Figure SSA).**

**Figure 7:**
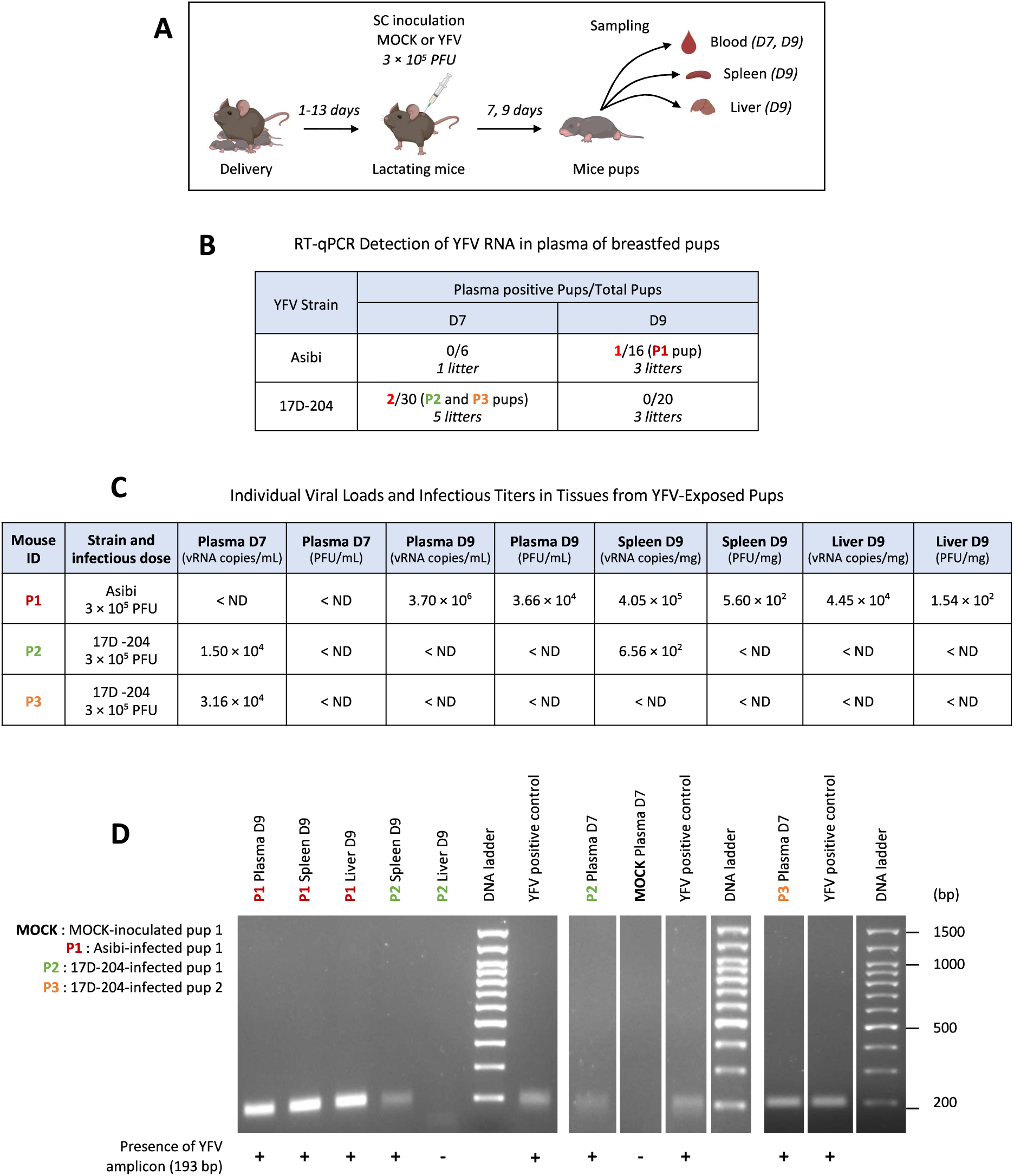
Limited but detectable transmission of YFV to suckling pups through breastfeeding. **(A)** Lactating A129 mice (12-24 weeks old) were inoculated subcutaneously (SC) 1 to 13 days postpartum with either PBS (MOCK, n=4) or 3 x 10^5^ PFU of Asibi (n=3) or 17D-204 (n=5) strains of YFV (3 independent experiments). Blood, spleen, and liver samples were collected from suckling pups at 7 (D7) and/or 9 (D9) days post-inoculation to assess transmission and viral dissemination (Asibi n=16, 3 litters; 17D-204 n =20, 5 litters) (Illustration created with Biorender). **(B)** Viral RNA levels in pup plasma were quantified by NS3-specific RT-qPCR. The same pups were analyzed at both time points: the 6 Asibi­ inoculated pups sampled at 7 dpi are part of the 16 analyzed at 9 dpi, and the 20 pups inoculated with 17D-204 at 9 dpi are included in the 30 analyzed at 7 dpi. ND (Not Detected) indicates no detectable RNA; NT (Not Tested) indicates that the condition was not included in the experimental design. **(C)** For plasma-positive pups, viral RNA and infectious titers were assessed in plasma, spleen, and liver by RT-qPCR and plaque assay (organs were homogenized prior to analysis). **(D)** In pups with plasma positive for YFV (C), the presence of viral RNA in spleen and liver was further evaluated by RT-qPCR following tissue homogenization. RT-qPCR-positive samples were then subjected to electrophoresis on a 2% TAE agarose gel to confirm amplification of the expected viral amplicons. A 100 bp Plus DNA ladder was used to estimate fragment size, with the expected amplicon corresponding to 193 base pairs (bp). Negative control included qPCR products from MOCK-infected pup samples (MOCK), while a plasmid containing YFV sequence (equivalent to 4 vRNA copies/µL) served as a positive control (YFV positive control). Full-size agarose gel electrophoresis images are shown in Supplementary Figure S4. MOCK: MOCK-inoculated pup 1; P1 : Asibi-infected pup 1; P2: 17D-204-infected pup 1; P3: 17D-204-infected pup 2.

Among all pups, three were tested positive for YFV RNA in plasma by RT-qPCR-1/16 fed by an Asibi-infected dam (P1) and 2/30 by 17D-204-infected dams (P2, P3) **(Figure 7B),** demonstrating that YFV breastfeeding transmission can occur, albeit infrequently.

Infected pups were further analyzed on plasma, spleen, and liver by RT-qPCR and plaque assay **(Figure 7C),** with confirmation of RT-qPCR amplicons by gel electrophoresis **(Figure 7D).** P1 showed high viral loads and infectious titers in all tissues at 9 dpi, with detection of the YFV-specific 193 bp amplicon, confirming systemic infection via breastfeeding. P2 and P3 had lower plasma RNA levels at 7 dpi; and at 9 dpi, only P2 showed low vRNA in the spleen and a positive viral amplicon.

These results demonstrate that both wild-type and vaccine YFV strains can be transmitted through breastfeeding in mice, although such transmission appears to be a rare event.

## DISCUSSION

Despite the availability of an effective live-attenuated vaccine, YFV remains a major public health concern. While mosquito transmission is primary, cases of vaccine-strain YFV transmission via breastfeeding, leading to meningoencephalitis in infants9-^12^, have prompted updated recommendations^13^• Detection of wild-type YFV RNA in human breast milk during outbreaks further raised concerns about potential non-vectorial transmission^14^• However, confirming this risk in endemic settings is difficult due to constant mosquito exposure. To address this, we conducted the first proof-of-concept study using an experimental animal model to investigate YFV transmission through breastfeeding and characterize underlying mechanisms.

We showed that both wild-type and vaccine YFV strains reach mammary glands in mice and are shed in breast milk at similar titers, though the vaccine strain disseminated to fewer glands. At 5-6 dpi, infectious virus was detected in 38.5% (Asibi) and 23.1% (17D- 204) of lactating mice - less frequently than in non-lactating mice (82.8% and 42.9%, respectively), possibly due to milk presence and physiological changes affecting detection.

Infectious virus of both YFV strains was detected in breast milk, whey and cell fraction, indicating the presence of free viral particles and cell-associated infectious virus. Despite reduced mammary dissemination, the vaccine strain reaches breast milk at levels similar to the wild-type strain, suggesting different mammary crossing mechanisms.

*In vivo,* YFV primarily targeted mammary stromal and immune cells, however mammary epithelial cells are also permissive *in vitro.* These findings suggest two potential routes into milk: direct epithelial infection with viral release or shedding, and infected immune cells transmigration ("Trojan horse strategy")-though other mechanisms cannot be excluded. Elucidating these mechanisms remains challenging and poorly defined, even for established breastfeeding-transmitted viruses like HIV, HTLV-1, and CMV^17^• Organoid models may help clarify epithelial susceptibility but may not fully recapitulate dynamics required to study viral crossing mechanisms^22^•

Crucially, both strains infected pups after oral exposure, with higher rates for Asibi (29.3%) than 17D-204 (8.5-9.7%), highlighting oral YFV transmission. Neuroinvasion occurred with both strains, especially Asibi (64.7% of infected pups). Together with reported neurological symptoms in cases of breastfeeding transmission of vaccine strains in humans^9–12^ Our findings raise concerns about the neurovirulence of wild-type YFV following oral exposure.

Using a human intestinal epithelium model, we found that differentiated enterocytes support productive viral replication, and apical-to-basal viral crossing *in vitro.* While transcytosis cannot be ruled out, paracellular passage seems unlikely given preserved barrier integrity.

Finally, as proof of concept, we confirmed rare but detectable YFV transmission through breastfeeding to suckling pups *in vivo*.

Overall, our findings highlight phenotypic differences between wild-type and vaccine YFV strains in dissemination and transmission efficiency. Both reach the mammary glands and are shed in milk at similar titers, despite less frequent gland infection with the vaccine strain. Oral infection experiments suggest higher neonatal susceptibility to the wild-type strain, indicating strain-specific differences in epithelial barriers crossing. These differences may be linked to the 20 amino acid differences-mainly in the envelope protein-between Asibi and 17D4, which may influence entry pathways and barrier crossing mechanism.

Previous work from our laboratory showed that ZIKV, another *Orthoflavivirus,* is transmitted via breastfeeding in A129 mice with markedly higher efficiency (39-90%)^18^•^23^ than YFV (∼6%). ZIKV also reached substantially higher milk titers (up to 10^9^ vs. 1.5 x 10^5^ PFU/ml)^18^ and showed greater oral infectivity (64% vs. 8.5% and 29.3% for vaccine and wild-type YFV)^23^. While interspecies differences must be considered, these findings suggest a markedly greater lactogenic transmission potential for ZIKV and underscore the relevance of the A129 model for comparing maternal flavivirus transmission.

Breastfeeding remains essential for infant health and is still recommended in ZIKV- endemic areas^24^ However, given confirmed vaccine-strain YFV transmission via breastfeeding in humans^9-12^, our findings reinforce the need to include this route among potential transmission risks for both YFV and ZIKV and to reassess current public health recommendations, especially in the absence of clear guidelines during maternal infection.

## SUPPLEMENTARY DATA

Supplementary materials are available online at———. These materials, provided by the authors to support and enhance the reader’s understanding, have not been copyedited and remain the sole responsibility of the authors. For any questions or comments, please contact the corresponding author.

## AUTHOR CONTRIBUTIONS

Conceptualization, P.-E.C. and A.V.;

Methodology, J.P., S.D., A.C., H.L. and A.V.;

Formal analysis, J.P.;

Investigation, J.P., S.D., P.J., R.K., A.C. and A.V;

Validation, H.L., A.V. and P.-E.C;

Resources, A.V. and A.C.;

Writing-original draft preparation, J.P.;

Writing-review & editing, J.P., S.D., A.C., H.L, A.G., P.-E.C, and A.V;

Supervision, A.V. and P.-E.C.;

Visualization, J.P., P.-E.C. and A.V.;

Project administration, P.-E.C. and A.V;

Funding acquisition, P.-E.C. and A.V.;

All authors have read and agreed to the published version of the manuscript

## ACKOWLEDGMENTS

We thank the staff of the Institut Pasteur ABSL 3 animal facility and the animal caretakers for their assistance with mouse experiments and occasional help with sampling; Nolwenn Jouvenet (lnstitut Pasteur, Paris, France) for providing viral strains, protocols, and plasmids; Valerie Choumet (lnstitut Pasteur) for generously providing antibodies; and Pascal Campagne (Bioinformatics and Biostatistics HUB, lnstitut Pasteur) for his support in improving the statistical analyses. We created some images (experimental diagrams) using a Biorender’s paid plan (by lnstitut Pasteur, Paris).

