## supplementary figures for "*In vivo* risk assessment of yellow fever virus transmission through breastfeeding, and mechanistic insights"

*Supplementary material*

### SUPPLEMENTARY FIGURES AND LEGENDS

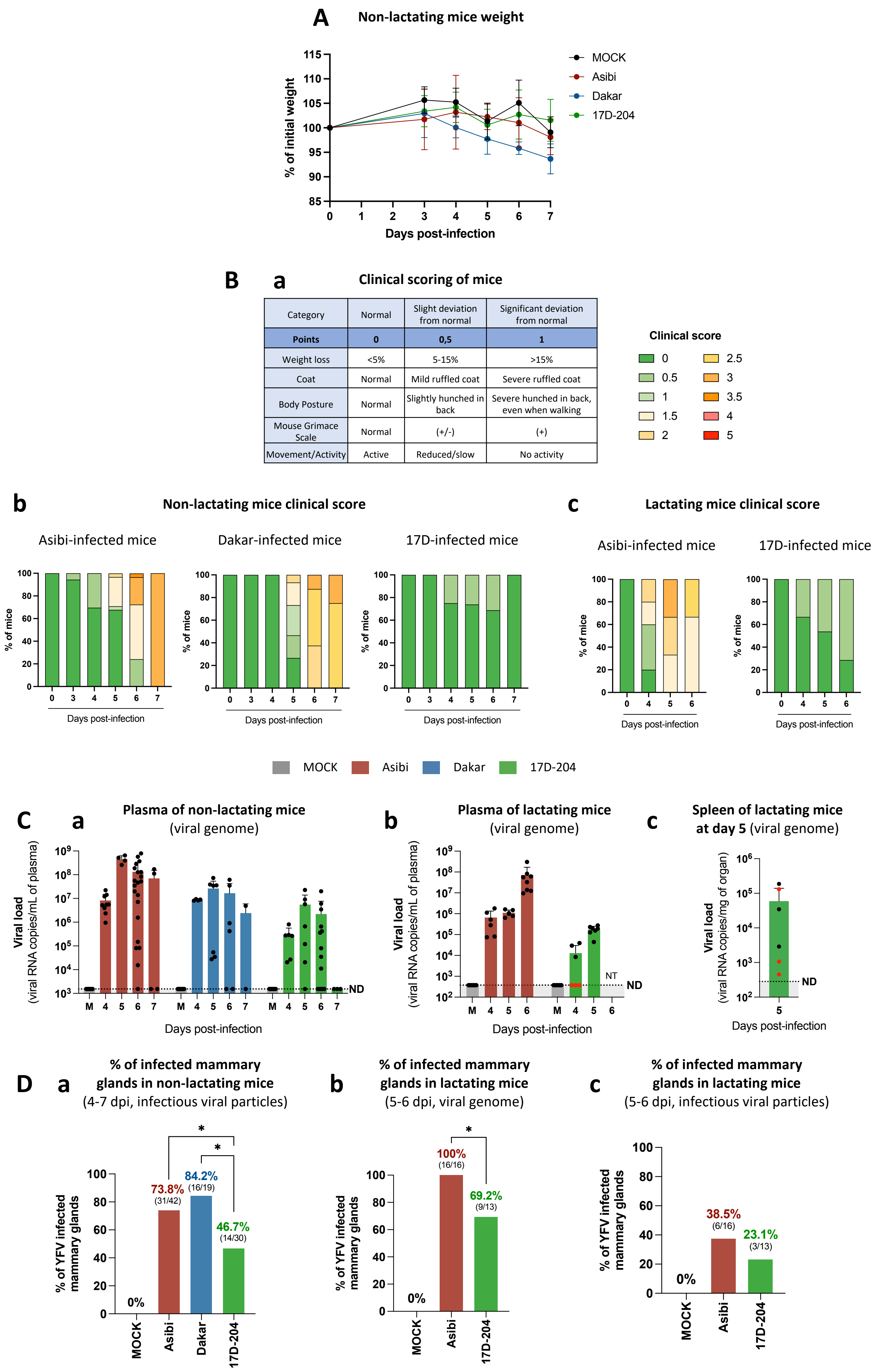


**Supplementary Figure S1 : clinical follow-up and validation of infection of non-lactating and lactating A129 mice.**
Non-lactating and lactating A129 mice were inoculated as described in Figure 1A. (**A-B**) Clinical monitoring was performed by assessing body weight in non-lactating mice (A), and by applying a standardized clinical scoring system (B). (**B**) Parameters included weight loss, coat condition, body posture, facial expression (Mouse Grimace Scale), and movement/activity (a). Each criterion was scored from 0 (normal) to 1 (severe deviation), with intermediate values reflecting slight deviations. The total clinical score ranged from 0 to 5. Mice reaching a cumulative score > 3.5 or showing marked deterioration in a single parameter were subjected to humane endpoint evaluation. See supplementary Materials and Methods for details. Clinical scores are shown for non-lactating (b) and lactating (c) mice. (**C**) Infection was confirmed by measuring viremia via RT-qPCR detection of viral RNA in plasma from non-lactating (a) and lactating mice (b). For lactating mice with undetectable plasma viremia at 4 dpi (red dots), infection was confirmed by detecting viral RNA in spleen tissue (c). (**D**) The proportion of infected mammary glands to the total number of mammary glands tested per condition was expressed as a percentage, for non-lactating mice (between 4 and 7 dpi) after detection by plaque assay (a), and for lactating mice (between 5 and 6 dpi), after detection by RT-qPCR (b) and plaque assay (c). Statistical tests: Fisher's exact test was used in panel (D). * = $p < 0.05$. Only statistically significant differences ($p < 0.05$) are indicated. Results in panels (A) and (C) are expressed as the mean ± standard deviation. The dashed lines in panel (C) indicate the specificity limit, which represents the threshold under which values were considered as "not detected" (ND). NT (Not Tested) indicates that the condition was not included in the experimental design.

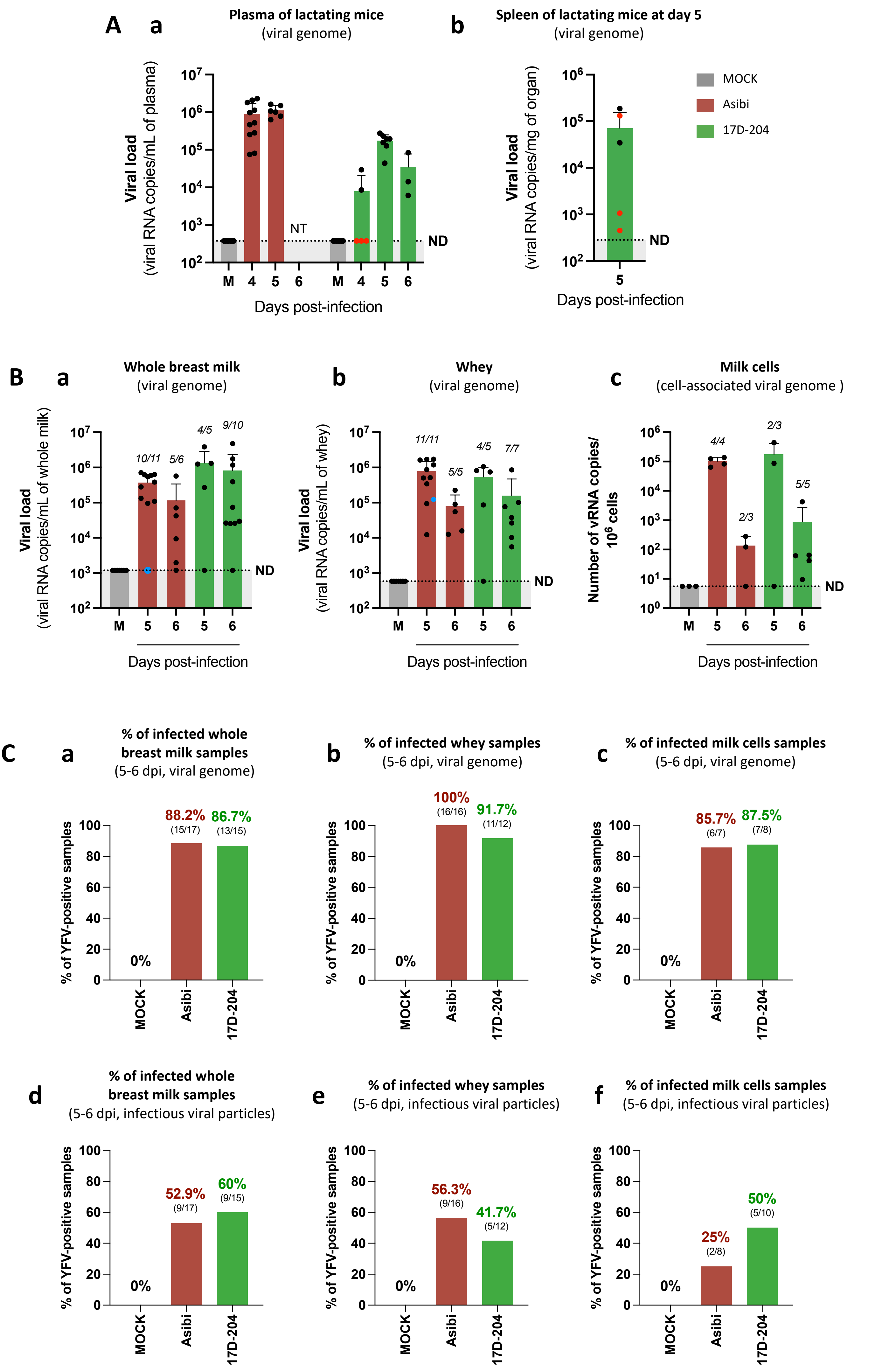


**Supplementary Figure S2 : Confirmation of YFV infection in lactating A129 mice and comparison of the proportion of infected milk samples between Asibi and 17D-204 strains.** Lactating A129 mice were inoculated as described in Figure 2A. (**A**) Infection was confirmed by measuring viremia via RT-qPCR detection of viral RNA in plasma from lactating mice (a). For lactating mice with undetectable plasma viremia at 4 days post-infection (dpi; red dots), infection was confirmed by detecting viral RNA in spleen tissue (b). NT (Not Tested) indicates that the condition was not included in the experimental design. Some data points from panel (a) and (b) are shared with panel S1C-b and S1C-c respectively, as they originate from the same mice used to collect both lactating mammary glands and maternal milk samples. (**B**) Detection of YFV in whole breast milk (**a**), whey (**b**) and milk cells (**c**) was assessed by NS3-specific RT-qPCR to detect viral RNA. Results are expressed as the mean ± standard deviation. The dashed lines indicate the specificity limit, which represents the threshold under which values were considered as "not detected" (ND). *The blue dot in panels (a) and (b) represents a whole milk sample with no detectable viral load but with a measurable infectious viral titer, along with the corresponding viral load and titer detected in the whey fraction isolated from this sample.* (**C**) The proportion of infected whole breast milk (a, d), whey (b, e) and milk cells (c, f) samples to total tested samples per condition was expressed as a percentage, for lactating mice (between 5 and 6 dpi) after detection by RT-qPCR (a-c) and plaque assay (d-f). Statistical tests: Kruskal–Wallis test with Dunn's multiple comparisons was used in panels (B) ; Fisher's exact test was used in panel (C). All comparisons indicated non-significant differences ($p > 0.366$).

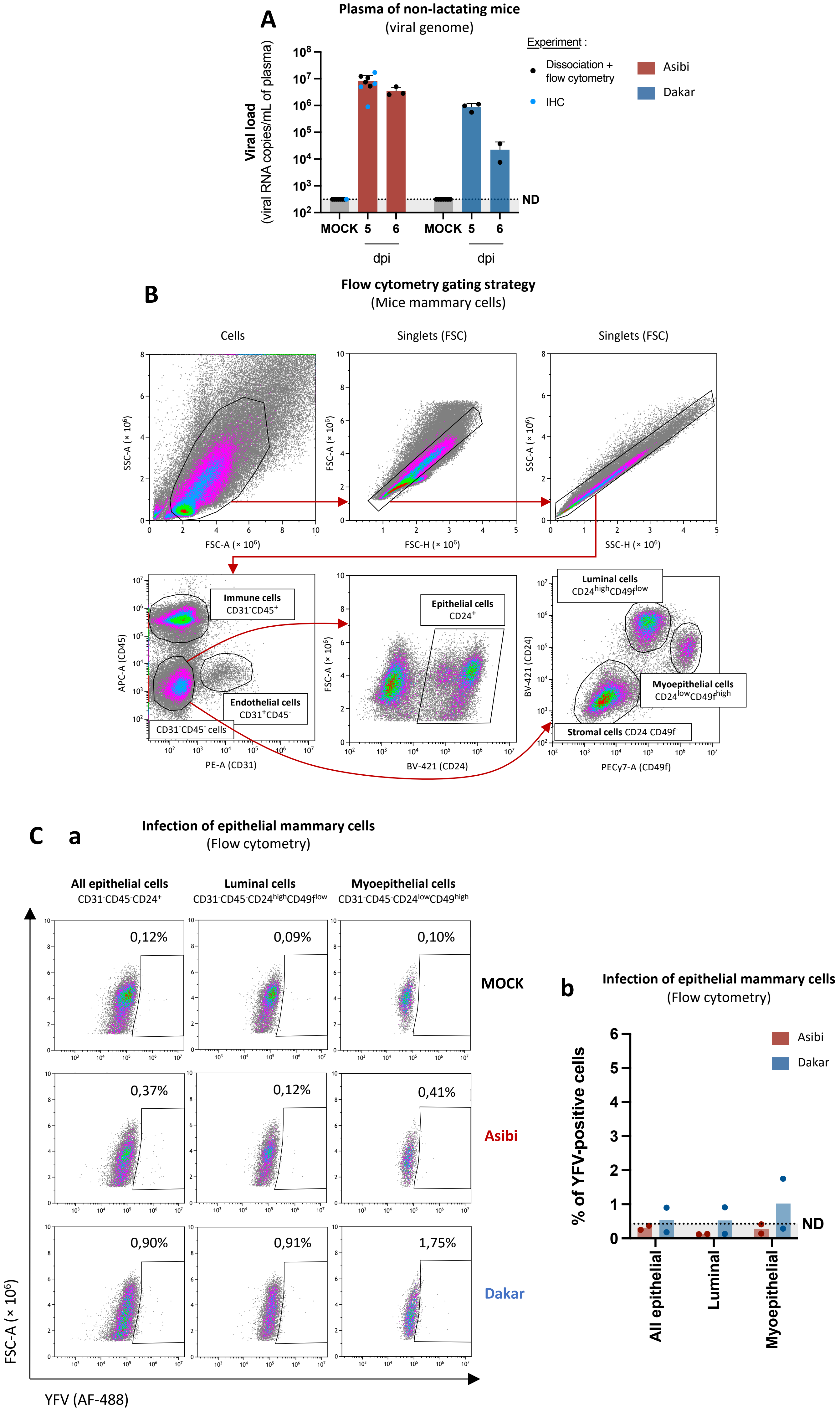


**Supplementary Figure S3 : Confirmation of YFV infection in A129 mice and cytometry gating strategy used for mice mammary cells.**
Non-lactating A129 mice were inoculated as described in Figure 3A. (**A**) At 5-6 days post-infection (dpi), infection was confirmed by measuring viremia via RT-qPCR detection of viral RNA. Results are expressed as the mean ± standard deviation. The dashed lines indicate the specificity limit, which represents the threshold under which values were considered as "not detected" (ND). (**B**) Representative flow cytometry gating strategy used to identify distinct mammary gland cell populations. Total events were first gated on forward scatter area (FSC-A) versus side scatter area (SSC-A) to exclude debris and select viable cells. Singlets were then gated using FSC height (FSC-H) versus FSC-A, and then SSC height (SSC-H) versus SSC-A to exclude doublets. From the singlet population, cells were first separated based on CD31 and CD45 expression: $CD31^+$ cells were classified as endothelial cells, and $CD45^+$ cells as immune cells. $CD31^-CD45^-$ cells were further analyzed for epithelial markers : total epithelial cells (all epithelial cells) were identified as $CD24^+$ cells, luminal epithelial cells were identified as $CD24^{high}CD49f^{low}$, while myoepithelial cells were defined as $CD24^{low}CD49f^{high}$. Stromal cells were defined as the quadruple-negative $CD31^-CD45^-CD24^-CD49f^-$ population. All populations were subsequently assessed for YFV infection as described in Figure 3B. (**C**)(**a**) Flow cytometry plots showing YFV-positive cells among all epithelial cells and luminal and myoepithelial cell subsets (1 representative experiment). (**b**) Quantification of YFV-positive epithelial cells (2 independent experiments, each dot represents one experiment). Results are presented as mean values; the dashed lines indicate the specificity limit, which represents the threshold under which values were considered as "not detected" (ND). These experiments correspond to those shown in Figure 3B.

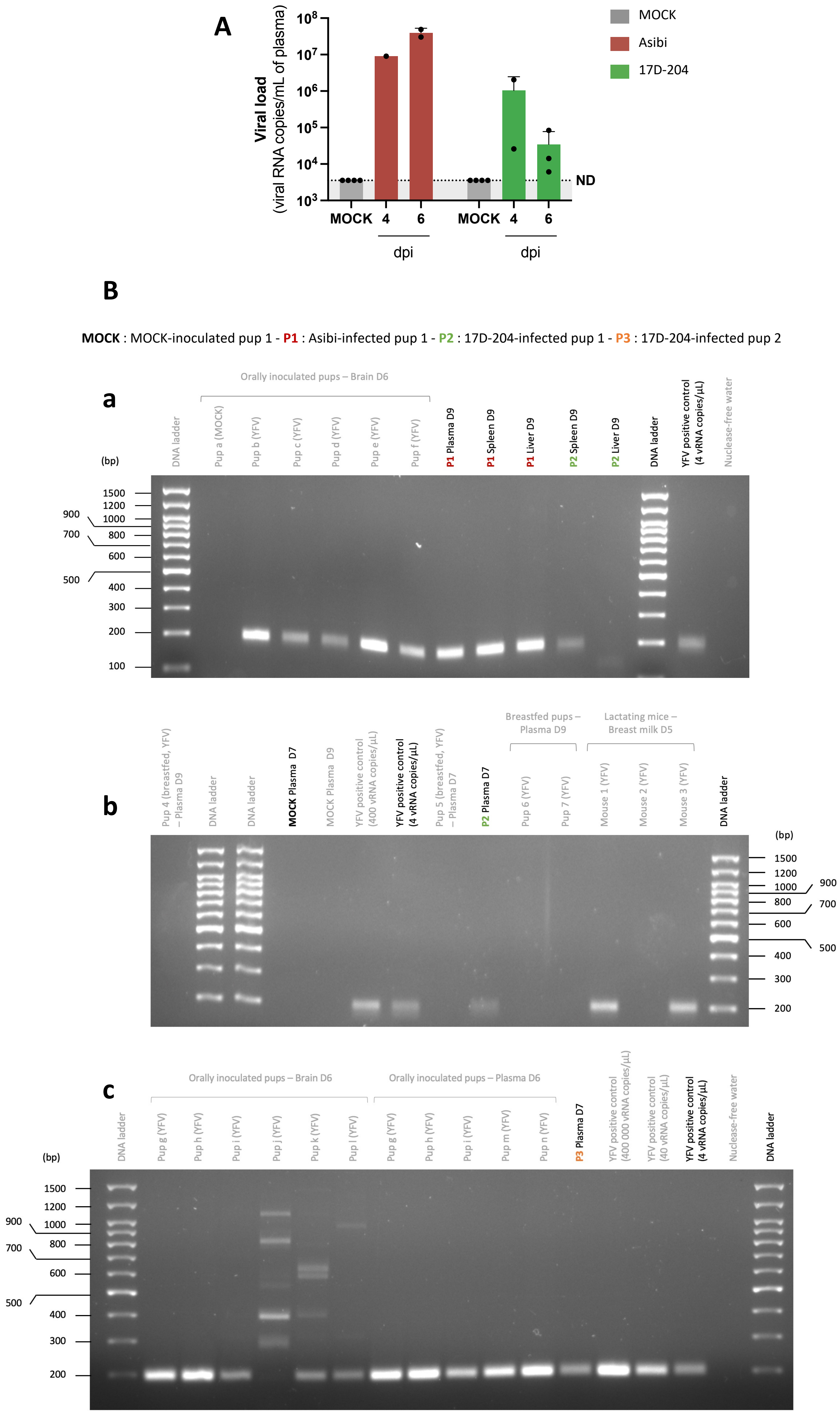


**Supplementary Figure S4: Confirmation of YFV infection in lactating A129 mice and validation of viral RNA in pups samples by full-size agarose gel electrophoresis.**
Lactating A129 mice were inoculated as described in Figure 8A. (**A**) Maternal infection was confirmed at 4 or 6 dpi by quantification of viral RNA in plasma via NS3-spectific RT-qPCR. Results are expressed as the mean ± standard deviation. The dashed lines indicate the specificity limit, which represents the threshold under which values were considered as "not detected" (ND). (**B**) Full-size agarose gel electrophoresis images corresponding to RT-qPCR amplicons shown in Figure 7D. RT-qPCR-positive samples, as well as additional samples from other experiments (indicated in grey), were subjected to electrophoresis on a 2% TAE agarose gel to confirm amplification of the expected viral amplicons. A 100 bp Plus DNA ladder was used to estimate fragment size, with the expected amplicon corresponding to 193 base pairs (bp). Negative controls included qPCR products from MOCK-infected pup samples (MOCK) and nuclease-free water, while a plasmid containing the YFV sequence at different concentrations (ranging from 4 to 400,000 vRNA copies/µL) served as positive controls (YFV positive controls). Gels are shown uncropped to display the full migration pattern of all bands. MOCK : MOCK-inoculated pup 1 ; P1 : Asibi-infected pup 1 ; P2 : 17D-204-infected pup 1 ; P3 : 17D-204-infected pup 2.
