## supplementary material and methods for "*In vivo* risk assessment of yellow fever virus transmission through breastfeeding, and mechanistic insights"

### MATERIALS AND METHODS

#### Virus strains

The three YFV strains (Asibi, Dakar/HD1279 and 17D-204) were kindly provided by N. Jouvenet (Institut Pasteur, Paris). Strains were amplified on Vero E6 cells and titrated by plaque assay as previously described[1], with minor modifications. Supernatants were collected at 72 h post-inoculation, clarified (3000 × g, 5 min), aliquoted, and stored at −80 °C for use in animal and *in vitro* experiments.

#### Cells

MDA-MB-231 cells (human mammary myoepithelial cell line, HTB-26, ATCC), Caco-2/TC7 cells (human enterocyte-like cell line, clone TC7, SCC209; Sigma-Aldrich, St. Louis, MO, USA), and Vero E6 cells (African green monkey kidney epithelial cell line, CRL-1586, ATCC) were grown in Dulbecco's Modified Eagle Medium (DMEM) supplemented with GlutaMAX (Gibco, Thermo Fisher Scientific, Waltham, MA, USA), 10% heat-inactivated fetal bovine serum (FBS) (Gibco), 100 U/mL penicillin and 100 µg/mL streptomycin (Gibco). MCF-7 cells (human mammary luminal cell line, HTB-22, ATCC) were grown in Dulbecco's Modified Eagle Medium F12 (DMEM/F12) supplemented with GlutaMAX (Gibco), 10% FBS, 100 U/mL penicillin, 100 µg/mL streptomycin, 20 ng/mL human epidermal growth factor (hEGF) (Peprotech, Neuilly-Sur-Seine, France), 0.5 µg/mL hydrocortisone (Sigma-Aldrich), and 10 µg/mL insulin (Sigma-Aldrich). Human primary mammary epithelial cells (HMEpiC) (ScienCell Research Laboratories, Carlsbad, CA, USA) were cultured in Mammary Epithelial Cell Medium (MEpiCM) supplemented with Mammary Epithelial Cell Growth Supplement (MEpiCGS) and penicillin/streptomycin (ScienCell) on poly-L-lysine-coated plates. All cells were maintained in an atmosphere of 5% $CO_2$ at 37 °C for culture.

#### Virus strains

Experiments were carried out using three YFV strains: two wild-type strains, Asibi strain (GenBank: AY640589.1) and Dakar/HD1279 strain (GenBank: MN106242.1); and one live-attenuated 17D-204 vaccine strain (GenBank: MN708488.1). The strains were kindly provided by Nolwenn Jouvenet (Institut Pasteur, Paris, France). Strains were amplified and titrated by plaque assay on Vero E6 cells as previously described[1], with minor modifications; supernatants were harvested at 72h post-inoculation and and clarified by centrifugation at 3000 × g for 5 min. Aliquots were then stored at −80°C and used as inoculum for both animal infections and *in vitro* assays.

#### *In vitro* model of human intestinal epithelium and verification of monolayer tightness

Caco-2/TC7 cells ($3 \times 10^4$ cells) were seeded onto the upper chamber of Transwell inserts (12 mm diameter, 3 µm pore size, polycarbonate, Costar). Apical (upper chamber) and basolateral (lower chamber) media were replaced by fresh medium three times per week, and cells were maintained in culture for 14 to 29 days. To validate monolayer tightness, Caco-2 monolayers integrity was monitored by measuring their trans-epithelial electrical resistance (TEER; EVOM World Precise Instruments) as previously described[2]. Caco-2 monolayers exhibiting TEER > 250 $\Omega.cm^2$ were considered impermeable, according to the criteria established in the literature[3].

**Cell infections**

Human mammary epithelial cells (HMEpiC, MDA-MB-231 and MCF-7 cells) were inoculated with the three YFV strains (Asibi, Dakar/HD1279, and 17D-204 strains) at different multiplicities of infection (MOI), as indicated in each experiment. After 2h of adsorption, cells were washed in phosphate-buffered saline (PBS) and the medium was replaced by fresh medium: MEpiCM medium for HMEpiC cells, DMEM-2% FBS for MDA-MB-231 cells or DMEM/F12-2% FBS for MCF-7 cells. At different time points, viral production in supernatants was assessed using plaque forming assay.

Differentiated Caco-2 tight monolayers (cultured for 14 to 29 days on Transwell inserts, with TEER ≥ 250 $Ohm.cm^2$) were inoculated in the apical compartment with the three YFV strains at an MOI of 1. Following 2h of adsorption, apical compartment was washed with PBS, and the medium was replaced by fresh DMEM containing 2% FBS. Monolayer tightness was monitored during all the experiment by measuring TEER, and morphological integrity was visualized 48 hours post-infection through immunofluorescence after immunostaining of the tight junction protein zonula occludens-1 (ZO-1). At various time points, infection was assessed by RT-qPCR after intracellular RNA extraction, and viral production in both apical and basolateral supernatants was measured using a plaque forming assay.

**Ethics statement**

Anonymized normal human primary mammary epithelial cells (HMEpiC) were obtained in compliance with local, state, and federal laws and regulations governing the procurement and distribution of human tissue, and provided by ScienCell Research Laboratories (Carlsbad, CA, USA).

**Mouse infection and sample collection**

A129 mice aged 6 to 24 weeks were used across different experiments. For non-lactating females, 6- to 23-week-old A129 animals were transferred to the ABSL-3 facility and subcutaneously inoculated with 1 to 3 × $10^5$ plaque-forming units (PFU) of YFV (Asibi, Dakar/HD1279 or 17D-204 strains). Blood samples were collected at various days post-inoculation (dpi) into tubes containing 15 mM EDTA (Invitrogen, Thermo Fisher Scientific), centrifuged at 2000 × g for 15 min, and the resulting plasma was stored at −80°C until further analysis by RT-qPCR. At different time points, mice were euthanized, and the abdominal and thoracic mammary glands were collected. Organs were either weighed and stored at -80 °C for plaque assays (thoracic glands), enzymatically dissociated for flow cytometry (abdominal and thoracic glands, with removal of inguinal lymph nodes for abdominal glands before analyses) or fixed for immunohistochemistry analyses (thoracic glands).

For experiments involving lactating females and offspring, 7- to 24-week-old A129 males and females were housed for breeding under approved animal welfare conditions. Following gestation and delivery, females were subcutaneously inoculated between 1 and 13 days post-partum with doses ranging from 1 × $10^5$ to 3 × $10^5$ PFU of the Asibi or 17D-204 strains of YFV. Blood samples from mice were collected between 4 and 6 dpi to monitor infection, and processed as previously described. For mammary gland collection, females and their litters were euthanized at 5 or 6 dpi, and thoracic and abdominal glands were harvested (inguinal lymph nodes removed for abdominal glands). Spleens were also collected, weighed, and stored at −80 °C for RT-qPCR analysis. For breast milk collection, samples were collected at 5 or 6 dpi as previously described[1], followed by euthanasia of dams and litters. For experiments of breastfeeding-mediated transmission, blood, liver, and spleen samples were collected from suckling pups at 7 (blood) and 9 (blood and organs) dpi, and analyzed as previously described. For oral infection of neonatal mice, following gestation and delivery, 4- to 7-days-old A129 pups were inoculated with doses ranging from $2 \times 10^4$ to $1 \times 10^5$ PFU of Asibi or 17D-204 strains of YFV by intragastric route using flexible gastric tubes. Blood samples and brains were collected at 6 dpi and analyzed as previously described.

Throughout all experiments, animals were monitored daily using a clinical scoring system based on body weight loss, coat condition, posture, activity, and facial expression (Mouse Grimace Scale). Scores were assigned as follows: 0 (normal), 0.5 (mild change), or 1 (marked change). Humane endpoints were strictly applied in accordance with ethical guidelines (see Supplementary Figure S1B).

**Breast milk fraction separation**

Whole milk samples were centrifuged at 1700 × g for 20min, to separate the cream, whey, and cellular pellet. The whey fraction was subsequently centrifuged three times at 5000 × g for 15min to remove residual cream. The cellular fraction was washed and centrifuged three times at 700 × g for 5min in PBS.

**Mammary gland dissociation and flow cytometry**

Freshly harvested thoracic and abdominal mammary glands from 3–4 mice per condition were pooled and placed in $CO_2$-independent medium (Gibco #18045-054), and mechanically minced using scissors and scalpels to obtain small tissue fragments. The fragments were incubated in $CO_2$-independent medium containing collagenase A (3 mg/mL, Roche #10103586001), hyaluronidase (100 U/mL, Sigma #H3884) and completed with 5% FBS for 1h30 at 37 °C under agitation (140 rpm). Following enzymatic digestion, cells were sequentially dissociated in pre-warmed 0.25% trypsin (Dutscher #P10-022100) and 0.1% EDTA Versen (Biochrom #L2113) for 1 min at room temperature (RT) followed by a wash in $CO_2$-independent medium + 5% FBS, then in Dispase II (5 mg/mL, Roche #13752000) supplemented with DNase I (0.1 mg/mL, Sigma Aldrich #D4527-40KU) for 5 min at 37 °C and finally in cold ammonium chloride solution (Stem Cell Technologies #07800) to lyse red blood cells. The resulting cell suspension was filtered through a 40 µm cell strainer. All the enzymes were diluted in complete $CO_2$-independent medium, and cells were centrifuged and washed in $CO_2$-independent medium between each dissociation step.

Mammary dissociated cells were immunostained with directly fluorochrome-conjugated antibodies against surface markers : CD31-PE (BD Biosciences # 561073, 2 µg/mL final), CD45-APC (clone 30-F11, Biolegend, 2 µg/mL final), CD24-BV421 (clone M1/69, Biolegend, 4 µg/mL final) and CD49f-PECy7 (a6-PE.Cy7, clone GoH3, Biolegend, 4 µg/mL final), diluted in $CO_2$-independent medium for 20min at 4°C in the dark. The cells were then fixed in PBS-4% paraformaldehyde (PFA) (Electron Microscopy Sciences, Hatfield,

PA, USA) for 15 min at RT, followed by permeabilization in PBS-0.1% Triton X-100 (Thermo Fisher Scientific) for 3 min at RT. YFV was labeled using YFV-specific mouse polyclonal antibodies (YFV ascitic fluid, 1/500, generously provided by Valérie Choumet, Institut Pasteur, Paris, France), diluted in PBS-0.2% Tween 20 (Sigma-Aldrich)-0.2% bovine serum albumin (BSA) (Sigma-Aldrich), and the incubation was carried out 30min at RT in the dark. A secondary Alexa Fluor 488-conjugated goat anti-mouse antibody (Invitrogen, 8 μg/mL final), diluted in PBS-0.2% BSA, was then applied for 30 min at RT in the dark. Cells were rinsed twice in PBS-0.1% BSA for 6 min at 700 × g between each step. Data were acquired via fluorescence-activated cell sorting (FACS) using CytoFLEX flow cytometer (Beckman Coulter, USA), and analysis was conducted using Kaluza (version 2.2, Beckman Coulter).

**Immunohistochemistry: tissue paraffin embedding, sectioning and immunostaining**
Freshly harvested thoracic mammary glands were fixed overnight (O/N) at RT with gentle agitation in 10% neutral-buffered formalin (Sigma Aldrich Chimie #HT501128-4L). Fixed tissues were dehydrated through a graded ethanol series (70%, 90%, and 100% (x2) ethanol, 45 min each) under agitation, followed by two 45-minute incubations in xylene. After clearing, samples were incubated for 1 hour at 60 °C in a 1:1 mixture of xylene and paraffin, then for 1 hour in pure paraffin at 60 °C, and finally in paraffin O/N. Tissues were embedded in paraffin using a Leica EG1150C embedding station (Leica Microsystems, Wetzlar, Germany), and stored at 4 °C. Sections of 5 μm thickness were cut using a microtome (Reichert-Jung 2030 Biocut manual microtome, Leica Microsystems) on cooled blocks, mounted on glass slides, and dried O/N at 37 °C. Slides were deparaffinized with two 5-minute xylene washes, then rehydrated through a descending ethanol series (100%, 90%, 70%, 50% ethanol) before rinsing in water. Heat-induced epitope retrieval (HIER) was performed by incubating slides in EDTA buffer (pH 8, Abcam #ab64216) at 121 °C for 20 minutes in a pressure cooker (Electron Microscopy Sciences #2100). Slides were then cooled to room temperature and rinsed with water.

Sections were processed for immunostaining in a humidified chamber. After two 5-min washes in Tris-buffered saline (TBS) containing 0.025% Triton X-100, non-specific binding was blocked by incubating the sections for 1h at room temperature in TBS-10% normal goat serum (NGS)-1% BSA. Sections were then incubated O/N at 4 °C with YFV-specific mouse polyclonal antibodies (YFV ascitic fluid, 1:500). The next day, sections were incubated for 1h at RT in the dark with the secondary antibody Alexa Fluor 546-conjugated goat anti-mouse IgG (Invitrogen, final concentration 8 μg/mL). Nuclei were then stained with DAPI (Abcam, #ab228549, final concentration 1 μg/mL) for 10 to 15 min at room temperature. Slides were washed and mounted using Immu-Mount (Epredia, Thermo Fisher Scientific). Images were acquired using an Olympus IX83 inverted fluorescence microscope (Olympus Corporation, Tokyo, Japan), and analyzed with ImageJ software. All antibodies and DAPI were diluted in TBS–1% BSA. Each incubation step was followed by two or three washes in TBS–0.025% Triton X-100, and in TBS after DAPI staining.

**RNA Extraction**
Mice organs (mammary glands, spleens, brains, and livers) were disrupted in TRIzol reagent (Ambion, Thermo Fisher Scientific) using a Bullet Blender Storm Homogenizer (Next Advance, Troy, MO, USA) with 3.2 mm stainless-steel beads (Next Advance) for 10

to 15 min at speed 12. Chloroform (Sigma-Aldrich) was added at 20% of the TRIzol volume, samples were vortexed and incubated for 10 min at RT, followed by centrifugation at 12000 × g for 20 min at 4°C to separate the different phases. The clear upper aqueous layer, containing RNA, was collected and incubated with cold isopropanol (VWR Chemicals, Radnor, PA, USA) for 1h at -20°C. After centrifuging at 12000 × g for 20 min at 4°C, the RNA pellet was washed twice with cold 70% ethanol (Thermo Fisher Scientific) and resuspended in nuclease-free water (Qiagen, Hilden, Germany). RNAs from plasma, whole milk, whey and cell supernatant samples were extracted using the QIAamp Viral RNA Mini Kit (Qiagen), according to the manufacturer's guidelines. RNA extractions from cell samples (mouse breast milk cells and Caco-2 cells) were carried out with the RNeasy Plus Mini Kit (Qiagen), according to the manufacturer's guidelines.

**RT-qPCR**

Reverse transcription was carried out using random hexamers and the Maxima H Minus Reverse Transcriptase kit (Thermo Fisher Scientific). Quantitative PCR was then performed with 5 µL of template cDNA, 10 µL of iTaq Universal SYBR Green Supermix (Bio-Rad, Hercules, CA, USA), and 500 nM of each YFV NS3-specific primer (Forward: 5'-GCG TAA GGC TGG AAA GAG TG-3'; Reverse: 5'-CTT CCT CCC TTC ATC CAC AA-3'). The PCR program was set up as follows: one cycle of 10 min at 95°C for initial denaturation, followed by 40 cycles of 15s at 95°C for denaturation, 20s at 60°C for annealing, and 30s at 72°C for extension. To assess the amplification specificity, a melting curve analysis was performed: the temperature was increased from 65 °C to 95 °C, in increments of 0.5 °C for 5s. Quantification was performed using a standard curve generated from a YFV-encoding plasmid. Experiments were carried out using the Eppendorf Realplex2 Mastercycler Epgradient S system (Eppendorf, Hamburg, Germany).

**Agarose gel electrophoresis**

A 2% agarose gel was prepared in 1X TAE buffer (40 mM Tris-acetate, 1 mM EDTA) containing ethidium bromide (Eurobio Scientific, Les Ulis, France). After solidification at RT for 30 min RT-qPCR products were mixed with 2 µL of 10X BlueJuice Gel Loading Buffer (Invitrogen) and 5 µL of the amplified DNA. Samples were loaded into the gel wells alongside a 100 bp Plus DNA Ladder (TransGen Biotech, Beijing, China). Electrophoresis was performed at 110 V for approximately 30 minutes. DNA bands were visualized under UV light using an Infinity gel system (Vilber Lourmat, Collégien, France).

**Viral titration by plaque forming assays**

Mice organs (mammary glands, spleens, brains, and livers) previously stored at -80 °C were thawed in PBS and dissociated using the Bullet Blender Storm Homogenizer with 3.2 mm stainless-steel beads for 4 min at speed 6. The suspensions were centrifuged at 2000 × g for 10 min, and the supernatants were collected for viral titration. Confluent monolayers of Vero cell were exposed to 10-fold dilutions of organ supernatant, plasma, whole milk, whey and cell supernatant samples in DMEM-2% FBS for 2 h at 37°C, 5% $CO_2$. After viral adsorption, the inocula were removed and replaced with DMEM-2% FBS-2% carboxymethyl cellulose (VWR). The cells were cultured for 7 days at 37°C, 5% $CO_2$. Following two washes in PBS, Vero monolayers were fixed with PBS-4% PFA for 15 min at RT and stained with crystal violet (Sigma-Aldrich).

#### Infectious center assays for milk cells

After 3-fold dilutions of 5 × $10^4$ to 5 × $10^5$ purified milk cells in DMEM containing 2% FBS, the cells were co-cultured with susceptible cells (Vero E6) for 2 hours at 37°C, 5% $CO_2$. After addition of DMEM-2FBS-2% carboxymethyl cellulose, the co-cultures were maintained for 7 days at 37°C, 5% $CO_2$. Following three washes in PBS, Vero monolayers were fixed in PBS-4% PFA for 15 min at RT and stained with crystal violet for 30 min.

#### Immunofluorescence

To assess YFV cell infection, Caco-2 cultured on Transwell inserts were rinsed with PBS and fixed in PBS-4% PFA for 15 min at RT. After permeabilization in PBS-0.5% Triton X-100 for 15 min at RT, non-specific binding sites were blocked using PBS-0.05% Tween 20-5% BSA for 30 min at RT. Tight junction ZO-1 protein were stained using a primary rabbit anti-ZO-1 antibody (Invitrogen, 2,5 µg/mL final) incubated for 2h at RT, and a secondary Alexa Fluor 546-conjugated donkey anti-rabbit antibody (Invitrogen, 4 µg/mL final) incubated for 1h at RT. Cells were rinsed twice with PBS between each step. Image acquisition was performed using a fluorescence microscope (EVOS FL, Life Technologies, Thermo Fisher Scientific), and image analysis was conducted using ImageJ version 1.54p software.

#### Statistical analysis

Statistical analyses were conducted using GraphPad Prism (version 10.4.2, GraphPad Software). The number of independent experiments and the statistical tests applied are specified in figure legends. Data are presented as mean values or mean ± standard deviation (SD). These analyses were conducted with guidance from engineers at the Bioinformatics and Biostatistics Hub of Institut Pasteur.
